# IFI16 senses and protects stalled replication forks

**DOI:** 10.1101/2025.04.07.646548

**Authors:** Amelia Gamble, Thomas A. Ward, Otto P.G. Wheeler, Caryl M. Jones, Laura G. Bennett, Ellen G. Vernon, Vithursha Thanendran, Jessica P. Morris, Ilaria Ceppi, Swagata Halder, Damiano Borello, Thomas D. J. Walker, Jahnavi Rajan, Gillian Dunphy, Petr Cejka, Leonie Unterholzner, Christopher J. Staples

## Abstract

Replication stress is a key driver of DNA damage and genome instability. Replication stress-induced fork remodelling generates a new DNA end that is vulnerable to the action of nucleases, and which is protected by a range of factors including the canonical tumour suppressors BRCA1 and BRCA2. Here we report that replication stress drives elevated production of cytokines and chemokines in the absence of DNA damage. The DNA sensor IFI16 binds nascent DNA at stalled replication forks and signals via the DNA sensing adaptor STING, to induce the activation of NF-κB and the production of pro-inflammatory cytokines in response to replication stress. IFI16 also acts directly at stalled replication forks to protect nascent DNA from degradation by the nucleases MRE11 and DNA2. Furthermore, IFI16 is required for the interferon-mediated rescue of fork protection in BRCA-deficient cells, highlighting the critical role of IFI16 in the cross-talk between innate immunity and fork protection during replication stress.

**Highlights:**

- Replication stress induces an early innate immune response, which is dependent on the DNA sensing factors IFI16 and STING, but not cGAS
- IFI16 binds directly to nascent DNA at stalled replication forks
- IFI16 prevents nucleolytic degradation of reversed forks
- IFI16 is required for interferon-mediated fork protection in BRCA-deficient cells

## Introduction

Replication stress arises consequent to impaired replication fork progression in response to a wide range of perturbations including nucleotide pool imbalance, polymerase inhibition, replisome encounter with the transcription machinery, or treatment with genotoxic agents^1^. Replication stress-induced DNA damage and genome instability are counteracted by a range of mechanisms that facilitate the preservation of nascent DNA integrity, the slowdown and resumption of DNA replication, and timely DNA repair^2^. The global physical remodelling (reversal) of replication forks by DNA translocases including SMARCAL1 and ZRANB3 is an important element of the replication stress response^8–10^. Fork reversal involves a template-switch that leads to the rehybridization of nascent DNA and the formation of a four-way DNA intermediate with a new and vulnerable end. Several canonical DNA repair factors (BRCA1, BRCA2, RAD51) act to protect these sites from the action of the nucleases MRE11, DNA2 and EXO1^3–7^.

DNA damage is also sensed by the cell-intrinsic innate immune system, leading to the production and secretion of type I interferons (IFN), cytokines, and chemokines^8^. Innate immune responses are driven by the recognition of pathogen- or damage-associated molecular patterns (PAMPs or DAMPs) by pattern recognition receptors (PRRs). Damaged DNA can act as a DAMP upon access to the cytosol, where it is detected by the DNA sensor cGAS (cyclic GMP-AMP synthase) and its adaptor protein STING (Stimulator of Interferon Genes)^9–11^. The innate immune system can also sense DNA damage in the nucleus. For instance, DNA damage induced by the topoisomerase II poison etoposide activates a STING-dependent, but cGAS-independent innate immune response in human cells^12^. This response involves nucleocytoplasmic shuttling of the DNA binding protein IFI16 (Interferon-γ-inducible factor 16), and the subsequent formation of a non-canonical STING signalling complex which induces a pro-inflammatory cytokine response *via* the activation of the transcription factor NF-κB^13^. However, it is still unclear which molecular DNA features are detected as DAMPs following nuclear DNA damage and replication stress.

Emerging evidence shows that there is a degree of interplay between components of the innate immune response and the replication stress response machinery. For instance, supplementation of BRCA1-deficient cells with IFNβ prevents MRE11-dependent nascent DNA degradation, though the underlying mechanism is currently unclear^14^. The innate immune DNA sensors cGAS and IFI16 bind DNA directly^15–18^, and have also been reported to exhibit STING-independent functions when bound to cellular or viral DNA. For instance, cGAS has been linked to the inhibition of DNA repair by homologous recombination, and the modulation of fork progression and nascent DNA integrity^19,20^. IFI16 has also been demonstrated to limit recruitment of DNA repair factors to sites of DNA damage^21^. Furthermore, IFI16 promotes the epigenetic modification and chromatinization of viral DNA, thus silencing viral gene expression independently of its function in innate immune signalling^22–24^. IFI16 can bind linear double-stranded (ds) DNA, as well as supercoiled and structured DNA, and has been proposed to scan along genomic DNA for stretches of DNA that lack nucleosomes^25^. However, it is currently unclear which DNA features are detected by IFI16 under conditions of replication stress and nuclear DNA damage.

Here, we demonstrate that replication stress initiates the rapid induction of an NF-κB-mediated innate immune response prior to elevated DNA damage or micronuclei formation. This response is driven by IFI16 and STING but is independent of cGAS. We show that IFI16 binds to nascent DNA at stalled replication forks, and that these structures are sensed as a danger signal during the acute inflammatory response to replication stress-inducing agents. We find that IFI16 also functions directly at stalled replication forks to prevent excessive resection of nascent DNA by the nucleases MRE11 and DNA2. Furthermore, we show that IFI16, as a highly IFN-inducible gene, contributes to the IFN-mediated rescue of nascent DNA degradation in BRCA1 or BRCA2-deficient cells. Conversely, IFI16 overexpression restores fork protection to BRCA-deficient cells independently of IFN treatment. Our data reveal a previously undiscovered function for IFI16 as a replication fork protection factor, and link fork stalling to activation of the inflammatory response during replication stress.

## Materials and Methods

### Cell culture and CRISPR-Cas9 cell line generation

U2OS, HaCaT cells, and Flp-In HEK293 cells were maintained as adherent monolayers in Dulbecco’s Modified Eagle’s medium (DMEM) with 10% Foetal Bovine Serum (FBS, Gibco). A375 melanoma cells, and T24 and J82 bladder cancer cells were grown in MEM (Gibco) containing 10 % FBS (Gibco). All cells were grown in 37°C in an atmosphere of 5% CO_2_. HaCaT cells lacking IFI16, cGAS or STING have been described previously^32^. Other IFI16 KO CRISPR clones were generated via lentiviral infection with Edit-R (Horizon) at an MOI of 0.3, followed by puromycin selection (Sigma, 2 μg/ml). Cells were reseeded into 96-well plates, and after a further 10 days, individual clones were subcultured and analysed by and Western blotting using an antibody raised against the IFI16 N-terminus to confirm deletion.

### Stable cell line generation

Stable HEK293 Flp-In cell lines expressing IFI16-FLAG were generated via cotransfection with pOG44 recombinase and either empty pDEST-Flag/FRT/TO or pDEST-FRT/TO-IFI16-FLAG according to the Flp-In manufacturer’s instructions (Invitrogen). Clones stably expressing IFI16-FLAG were selected in medium containing blasticidin S (15 μg/ml) and hygromycin B (200 μg/ml) (Invitrogen).

### RNAi transfections

Cells were transfected with 10-50 nM siRNA, depending on cell type using Lipofectamine RNAiMAX or Lipofectamine 2000 (Invitrogen) according to the manufacturer’s instructions. siRNAs used were pools of 4 siRNAs (ON-TARGET Plus SMARTpool siRNA, Dharmacon): non-targeting pool (Cat No: D-001810-10), IFI16 (L-020004), SMARCAL1 (L-013058), ZRANB3 (L-010025). Cells were subsequently further treated, lysed or fixed for analysis at least 48 hrs post-transfection unless otherwise indicated.

### Cell lysis and Western blotting

For the preparation of whole-cell extracts, cells were solubilized in lysis buffer [25 mM Tris-HCl (pH 7.4), 150 mM NaCl, 1% Triton X-100, 1 mM dithiothreitol (DTT), and 1 mM MgCl_2_] supplemented with Benzonase (50 U/ml) (Novagen) and cOmplete protease inhibitors and PhosSTOP phosphatase inhibitors (Roche). Lysates were clarified by centrifugation at 16,000*g* for 15 min at 4°C. Alternatively, cells were lysed directly in sample buffer (62.5mM Tris-Cl pH 6.8, 2% SDS, 10%g glycerol, 0.1% Bromophenol Blue, 50mM DTT). Gel electrophoresis was performed using 4-12% NuPAGE gels (Invitrogen) in MOPS running buffer (Invitrogen) or 12% polyacrylamide gels (BioRad Tetra system) in running buffer containing 0.25M Tris, 2M glycine and 1% SDS. Resolved proteins were transferred to polyvinylidene difluoride (PVDF) membranes, which were then probed for the protein of interest using antibodies diluted in phosphate-buffered saline (PBS)–0.1% Tween 20 (Sigma-Aldrich) containing 5% Marvel or 5% BSA (Sigma) as appropriate. Blots were imaged on a Bio-Rad ChemiDoc machine or an iBright imager (Invitrogen).

### Immunofluorescence

Cells were grown on glass coverslips in 24-well trays, transfected/treated as indicated, fixed with 3% PBS-buffered paraformaldehyde for 10 min at room temperature. For pre-extraction, cells were incubated with CSK-Trinton buffer (10mM PIPES pH6.8, 100mM NaCl, 300mM sucrose, 1.5 mM MgCl_2_, 5mM EDTA, 0.5% Triton-X100) for 4 min on ice, prior to fixation. Cells were permeabilized for 12 min in PBS containing 0.5% Triton X-100 for 5 min at room temperature, blocked in 3% BSA in PBS for 1h, and incubated with primary antibody in blocking buffer or PBS overnight in the cold room. Secondary antibody incubation was carried out for 3h at room temperature with Alexa Fluor 488– or Alexa Fluor 594–conjugated goat anti-rabbit or anti-mouse immunoglobulin G fluorescent secondary antibodies (Invitrogen, 1:1000) in blocking buffer or PBS. Washes after incubations were performed in PBS. DNA was counterstained with 4′,6-diamidino-2-phenylindole (DAPI, 1 μg/ml), and coverslips were mounted cell-side down in Shandon Immu-Mount medium (Thermo Fisher Scientific) or MOWIOL 4-88 (Sigma) containing 1 μg/ml DAPI. Fluorescence microscopy was performed on a Zeiss LSM710 or Zeiss LSM880 confocal microscope at ×40 or ×63 magnification. Images were captured and analysed using Zen software (Zeiss) and ImageJ. For the quantification of nuclear γH2A.X intensity using ImageJ, masks for individual nuclei were generated using the DAPI channel and used to quantify median intensity of γH2A.X staining in each nucleus. At least 50 nuclei were scored per sample.

### Clonogenic survival assays

Cells were plated onto 10 cm culture dishes and treated with genotoxic agents as required. After 24 hrs the media was changed. Colonies were incubated for 12 days, and then were fixed and stained using methylene blue/methanol. Colonies were counted manually, and results normalised to untreated controls.

### DNA fibre assay

Cells were plated and transfected appropriately, then were pulse-labelled with 25 μM CldU (Sigma-Aldrich) and 250 μM IdU (Sigma-Aldrich) as indicated in combination with replication stress-inducing agents and/or inhibitors as required. Cells were then harvested in PBS. Nuclei were then pelleted and resuspended in PBS. Two point five microlitres of suspended nuclei were mixed with 7.5 μl of lysis buffer (200 mM Tris pH 7.4, 25 mM EDTA, and 0.5% SDS) on a clean, dry slide (Thermo Fisher Scientific). After 7 min, slides were tilted at 30°, air-dried, then fixed in cold methanol/acetic acid (3:1). DNA fibres were denatured using 2.5 M HCl for 75 min then washed with 1× PBS before blocking in 1% Bovine Serum Albumin (BSA)/PBS containing 0.2% Tween 20 for 1 hour. CldU- and IdU-labeled tracts were incubated with two anti-BrdU (5-bromo-2′-deoxyuridine) antibodies, one of which is specific for CldU (Abcam) and the other for IdU (BD). Slides were then washed and incubated with goat anti-mouse/rat Alexa Fluor 488 and Alexa Fluor 594 (Invitrogen). DNA fibres (150 per condition) were visualized on a Zeiss LSM710 confocal microscope, and images were collected using Zen software and then analysed with ImageJ.

### iPOND

iPOND was performed on HeLa cells exactly as described previously by Sirbu and colleagues^26^. Briefly, newly synthesized DNA was labelled with 10 μM EdU for 10 min, cells were fixed in 1% formaldehyde and permeabilized, and the click reaction was performed using azide–polyethylene glycol biotin conjugate (Click Chemistry Tools). Cell preparations were then sonicated, and EdU-labelled DNA was precipitated using streptavidin beads, before washing and elution in loading buffer containing 1 mM DTT.

### Expression and purification of recombinant proteins

The *IFl16* sequence was amplified by PCR from the pCMV-HA-IFl16 plasmid (Origene) using the IFl16_Fw (CCTGCTGCTAGCATGGGAAAAAAATACAAGAACA) and IFl16_Rev (AGCAGGCCCGGGGTCTGGTGAAGTTTCCATACTT) primers containing the NheI and XmaI restriction sites, respectively. The *IFl16* gene was then cloned into pFB-2XMBP-HLTF-10Xhis^27^ to generate pFB-2XMBP-IFl16-10Xhis. IFl16 was expressed in *Spodoptera frugiperda* 9 (*Sf*9) cells in SFX Insect serum-free medium (Hyclone) using the Bac-to-Bac expression system (Invitrogen), according to manufacturer’s recommendations. Frozen *Sf*9 pellets from 0.6l culture were resuspended in lysis buffer (50 mM Tris-HCl pH 7.5, 1 mM EDTA, 1:300 protease inhibitor cocktail [Sigma], 30 µg/ml leupeptine [Merck Millipore], 1 mM PMSF, 5 mM β-mercaptoethanol) and incubated at 4°C for 20 min. Glycerol was added to a final concentration of 25%, NaCl was added to a final concentration of 305 mM and the solution was incubated at 4°C for 30 min. The cell suspension was centrifuged at 55,000 g at 4°C for 30 min. The soluble extract was incubated with amylose resin (New England Biolabs) at 4°C for 1 h. The resin was washed with amylose wash buffer 1 (50 mM Tris-HCl pH 7.5, 5 mM β-mercaptoethanol, 1 M NaCl, 10% glycerol, 1 mM PMSF), followed by one last wash with amylose wash buffer 2 (50 mM Tris-HCl pH 7.5, 5 mM β-mercaptoethanol, 300 mM NaCl, 10% glycerol, 1 mM PMSF). Proteins were eluted using amylose elution buffer (50 mM Tris-HCl pH 7.5, 5 mM β-mercaptoethanol, 300 mM NaCl, 10% glycerol, 1 mM PMSF, 10 mM maltose [Sigma]). The MBP-tagged variants were incubated with PreScission protease (20 mg PreScission protease per 100 mg of tagged protein) at 4°C for 1 h to cleave the MBP-tag. Subsequently, imidazole [Sigma] was added to a final concentration of 20 mM and the solution was incubated with pre-equilibrated Ni-NTA agarose resin (Qiagen) at 4°C for 1 h, in agitation. The resin was washed with Ni-NTA buffer 1 (50 mM Tris-HCl pH 7.5, 2 mM β-mercaptoethanol, 1 M NaCl, 10% glycerol, 1 mM PMSF, 20 mM imidazole) and subsequently with Ni-NTA buffer 2 (50 mM Tris-HCl pH 7.5, 2 mM β-mercaptoethanol, 150 mM NaCl, 10% glycerol, 1 mM PMSF, 20 mM imidazole). Proteins were eluted with Ni-NTA elution buffer (50 mM Tris-HCl pH 7.5, 2 mM β-mercaptoethanol, 150 mM NaCl, 10% glycerol, 1 mM PMSF, 200 mM imidazole). Fractions containing high protein concentration as estimated by Bradford were pooled and dialyzed at 4°C for 1 h in dialysis buffer (50 mM Tris-HCl pH 7.5, 2 mM β-mercaptoethanol, 100 mM NaCl, 10% glycerol, 0.5 mM PMSF). The dialyzed protein was aliquoted, snap-frozen in liquid nitrogen and stored at −80°C.

MRE11, WRN, DNA2 and RPA were expressed in *Sf*9 cells in SFX Insect serum-free medium (Hyclone) using the Bac-to-Bac expression system (Invitrogen), according to manufacturer’s recommendations. MRE11 was purified by affinity chromatography exploiting the N-terminal maltose-binding protein (MBP)-tag and the C-terminal polyhistidine (his)-tag^28^. The MBP-tag was removed before the NiNTA affinity purification step using PreScission Protease. WRN was purified exploiting the MBP-tag at the N-terminus and 10Xhis-tag at the C-terminus^29^. DNA2 was purified by affinity chromatography exploiting the N-terminal 6Xhis-tag and the C-terminal flag-tag^29^ RPA was purified using Ni-NTA affinity resin (Qiagen), followed by ÄKTA pure (GE Healthcare) with HiTrap Blue HP column (GE Healthcare), HiTrap desalting column (GE Healthcare) and HiTrap Q column (GE Healthcare)^30^.

### ELISAs

Cells were seeded in 96-well plates as triplicate samples and stimulated for 24h or as indicated. Supernatants were harvested, and the amount of secreted IL-6 protein was quantified using Human IL-6 DuoSet ELISA kits (DY206, R&D Systems) according to manufacturer’s instructions. Absorbance was measured at 450nm and corrected against absorbance at 570nm on a Tecan Infinite 200 Pro plate reader. Assays were carried out with triplicate samples and concentrations were calculated based on a standard curve with recombinant IL-6.

### qRT-PCR

Cells were seeded in 12-well plates as triplicate samples, and stimulated for 4-48h, as indicated. RNA was extracted using an E.Z.N.A Total RNA kit (Omega Bio-Tek) according to manufacturer’s instructions, including an in-column treatment strep with RNAse-free DNAse I (Omega Bio-Tek). cDNA synthesis was performed using the iScript Advanced cDNA Kit for RT-qPCR (Bio-Rad). qRT-PCR was carried out using Fast SYBR Green Master Mix (Applied Biosystems) on a BioRad CFX opus 96 instrument. Primers were designed to span at least one exon-exon junction. Ct values for mRNAs of interest were normalised to Ct values for GAPDH or β-actin mRNA, and data is expressed as fold change over mock treatment for control cells (wild type or scrambled siRNA). Primer sequences were: β-actin forward (FWD): 5’- CGCGAGAGAAGATGACCCAGATC-3’; β-actin reverse (REV): 5’- GCCAGAGGCGTACAGGGATA-3’; GAPDH FWD: 5’- CTCCTGTTCGACAGTCAGCC-3’; GAPDH REV: 5’- ACCAAATCCGTTGACTCCGAC-3’; IL6 FWD: 5’-CAGCCCTGAGAAAGGAGACAT- 3’, IL6 REV: 50-GGTTCAGGTTGTTTTCTGCCA-3’; CCL20 FWD: 5’- AACCATGTGCTGTACCAAGAGT-3’; CCL20 REV: 5’- AAGTTGCTTGCTTCTGATTCGC-3’; IFNβ FWD: 5’-ACGCCGCATTGACCATCTAT-3’; IFNβ REV: 5’-GTCTCATTCCAGCCAGTGCT-3’; CXCL10 FWD: 5’- AGCAGAGGAACCTCCAGTCT-3’; CXCL10 REV: 5’- AGGTACTCCTTGAATGCCACT-3’; For the qRT-PCR array, cDNA was prepared using RT2 First Strand Mix (QIAGEN), and amplified using RT2 SYBR Green qPCR Master Mix (QIAGEN) with the RT2 Profiler PCR Array Human Cytokines and Chemokines (PAHS-150ZF, QIAGEN), according to manufacturer’s instructions. Data was analysed using SABioscience PCR array analysis software.

### DNA substrate preparation

The DNA substrates for the *in vitro* assays were radiolabeled at the 3ꞌ terminus with [α-^32^P]dCTP (Hartmann Analytic) and terminal transferase (New England Biolabs) according to the manufacturer’s instructions. Unincorporated nucleotides were removed using Micro Bio-Spin P-30 Tris chromatography columns (Biorad). The 4-way dsDNA junctions substrate was prepared by annealing of the oligonucleotides X12-3TOPL (GACGTCATAGACGATTACATTGCTAGGACATGCTGTCTAGAGACTATCGCGACTTACGTTCCATCGCTAGGTTATTTTTTTTTTTTTTTTTTT), X12-3HJ3 (GAGATCTATCTGGTGCCTTCTGACAGTGAATGGGTAACGAATCGTAATAGTCTCTAGACAGCATGTCCTAGCAATGTAATCGTCTATGACGTC), X12-3HJ2 (CCCAAGTACGGTTGATTCGGGGCCAGTAGCATCCTAGTTAAGCCCATTACGATTCGTTACCCATTCACTGTCAGAAGGCACCAGATAGATCTC), and X12-3HJ1 (AAAAAAAAAAAAAAAAAAATAACCTAGCGATGGAACGTAAGTCGCGATGGGCTTAACTAGGATGCTACTGGCCCCGAATCAACCGTACTTGGG). The 3-way dsDNA DNA junction substrate was prepared by annealing of the oligonucleotides X12-3TOPL (GACGTCATAGACGATTACATTGCTAGGACATGCTGTCTAGAGACTATCGCGACTTACGTTCCATCGCTAGGTTATTTTTTTTTTTTTTTTTTT) and X12-3HJ3 (GAGATCTATCTGGTGCCTTCTGACAGTGAATGGGTAACGAATCGTAATAGTCTCTAGACAGCATGTCCTAGCAATGTAATCGTCTATGACGTC). The dsDNA substrate was prepared by annealing of the oligonucleotides X12-3HJ3 (GAGATCTATCTGGTGCCTTCTGACAGTGAATGGGTAACGAATCGTAATAGTCTCTAGACAGCATGTCCTAGCAATGTAATCGTCTATGACGTC) and X12-3HJ3-C (GACGTCATAGACGATTACATTGCTAGGACATGCTGTCTAGAGACTATTACGATTCGTTACCCATTCACTGTCAGAAGGCACCAGATAGATCTC). The 2.2 kbp-long dsDNA was prepared as published before^31^ by amplifying the human *NBS1* gene by PCR containing 66 nM [α-^32^P]dCTP together with the standard dNTPs concentration (200 µM each).

### Nuclease Assays

The exonuclease assays with recombinant MRE11 and IFI16 were carried out in a 15 μl volume. The reactions were assembled on ice in reaction buffer containing 25 mM Tris-acetate (pH 7.5), 1 mM DTT, 5 mM magnesium acetate, 3 mM manganese acetate, 1 mM adenosine triphosphate (ATP), 80 U/ml pyruvate kinase (Sigma-Aldrich), 1 mM phosphoenolpyruvate, 0.25 mg/ml BSA (New England Biolabs), and 5 nM ^32^P-labelled oligonucleotide-based dsDNA substrate (in molecules). The reactions were incubated for 2 h at 37°C and stopped by adding 0.5 μl of proteinase K (20.3 mg/ml, Roche) and 0.5 μl of 0.5 M EDTA for 30 min at 50°C. The stopped reactions were mixed with an equal volume of loading dye [95% formamide, 20 mM EDTA, and bromophenol blue (1 mg/ml)] and boiled for 4 min at 95°C. The reaction products were separated by denaturing electrophoresis on 15% denaturing polyacrylamide gels (acrylamide:bisacrylamide, 19:1; Bio-Rad) in 1x TBE buffer (89 mM tris, 89 mM boric acid, and 2 mM EDTA). The resolved gels were fixed in fixing solution (40% methanol, 10% acetic acid, and 5% glycerol) for 30 min at room temperature. The gels were then dried, exposed to storage phosphor screen, and scanned by a Typhoon imager (GE Healthcare). Quantitation was carried out using ImageJ/Fiji software.

The DNA end resection assays with PCR-based substrate were performed in 15 µl volume in a reaction buffer containing 25 mM Tris-acetate (pH 7.5), 2 mM magnesium acetate, 1 mM DTT, 0.1 mg/ml BSA (New England Biolabs), 1 mM phosphoenolpyruvate, 80 U/ml pyruvate kinase (Sigma), 50 mM NaCl, 1 nM ^32^P-labelled PCR-based DNA substrate (in molecules). The reaction was assembled on ice with the indicated proteins, and incubated at 37°C for 30 min. Next, 5 µl of 2% STOP solution (150 mM EDTA, 2% sodium dodecyl sulphate, 30% glycerol, 0.1% bromophenol blue) and 1 µl of Proteinase K (20.3 mg/ml, Roche) was added to stop the reaction for 10 min at 37°C. Samples were analysed by 1% agarose gel electrophoresis, dried on DE81 chromatography paper (Whatman), exposed to storage phosphor screens (GE Healthcare) and scanned by Typhoon FLA 9500 (GE Healthcare). Quantitation was carried out using ImageJ/Fiji software.

### EMSAs

DNA binding experiments (15 µl volume) with oligonucleotide-based DNA were carried out in buffer containing 30 mM Tris-HCl (pH 7.5), 1 mM DTT, 0.1 mg/ml BSA (New England Biolabs), 2 mM magnesium chloride, 1 mM ATP and 1 nM of the indicated ^32^P-labelled PCR-based DNA substrate (in molecules). The reaction was assembled on ice and incubated at 37°C for 15 min. After incubation, the samples were supplemented with 5 µl loading dye (50% glycerol with bromophenol blue) and analysed by gel electrophoresis on a 6% polyacrylamide gel in 1x TAE. Gels were run for 1 h at 100 V on ice, dried and exposed to a phosphor screen. The screen was then imaged using a Typhoon FLA 9000 (GE Healthcare).

### List of antibodies

The following antibodies were used.

*Western blotting*: GAPDH Antibody (SCBT, G-9, 1:2000), RPA2 (Abcam 9H8, ab2175, 1:1000), Tubulin (Abcam, ab4074), SMARCAL1 (SCBT, sc-376377), γH2AX (Cell Signaling Technology, 9718), PCNA (SCBT, PC10, sc-56, 1:1000), H3 (Cell Signaling Technology, 9715, 1:1000), IFI16 (SCBT, 1G7, sc-8023, 1:1000); cGAS (Cell Signaling Technology, E5V3W, 1:1000), STING (Cell Signaling Technology, D2P2F, 1:1000), β-actin (Sigma A2228, 1:3000).

*DNA Fibre Assays*: Anti-BrdU (BD 347580, 1:250; cross-reacts with IdU). Anti-BrdU BU1/75 (Abcam, ab6326, 1:250).

*Immunofluorescence*: 53BP1 (Abcam, ab21083, 1:2000), γH2AX (Upstate, 1:1000 or Cell Signaling Technology, 2577, 1:600), IFI16 (SCBT, 1G7, sc-8023 or Cell Signaling Technology D8B5T, 1:600), NF-κB p65 (Cell Signaling Technology 6956 or 8242, 1:300), IRF3 (Cell Signaling Technology 11904, 1:300).

## Results

### Replication stress induces a STING-dependent, but cGAS-independent pro-inflammatory cytokine response

We have previously shown that the genotoxic agent etoposide causes the non-canonical activation of STING in human epithelial cells, leading to NF-κB activation and pro-inflammatory cytokine production^12^. Etoposide promotes trapping of DNA loops by topoisomerase II, which generates a roadblock to DNA replication, and leads to DNA double-strand break (DSB) formation. To determine whether replication stress generates a proinflammatory cytokine response even in the absence of DNA damage, we treated human HaCaT keratinocytes with the replication stress inducer hydroxyurea (HU), which causes fork stalling by dNTP depletion. We observed that HU treatment leads to the nuclear translocation of NF-κB p65 as early as 2h post treatment but induces only minimal translocation of the transcription factor IRF3 (Fig. 1A-C). In contrast, transfection with dsDNA results in a cytosolic DNA sensing response which involves the predominant activation of IRF3, and only modest NF-κB p65 activation (Fig. 1A-C). HU-induced replication stress resulted in the increased expression of a range of pro-inflammatory cytokines and chemokines as quantified by qPCR array (Fig. 1D). The cytokine expression profile resembles that induced by etoposide treatment, and is distinct from the response to cytosolic DNA, which is characterised by the induction of chemokines such as CXCL10 and CCL5 (Fig. 1D).

**Figure 1:**
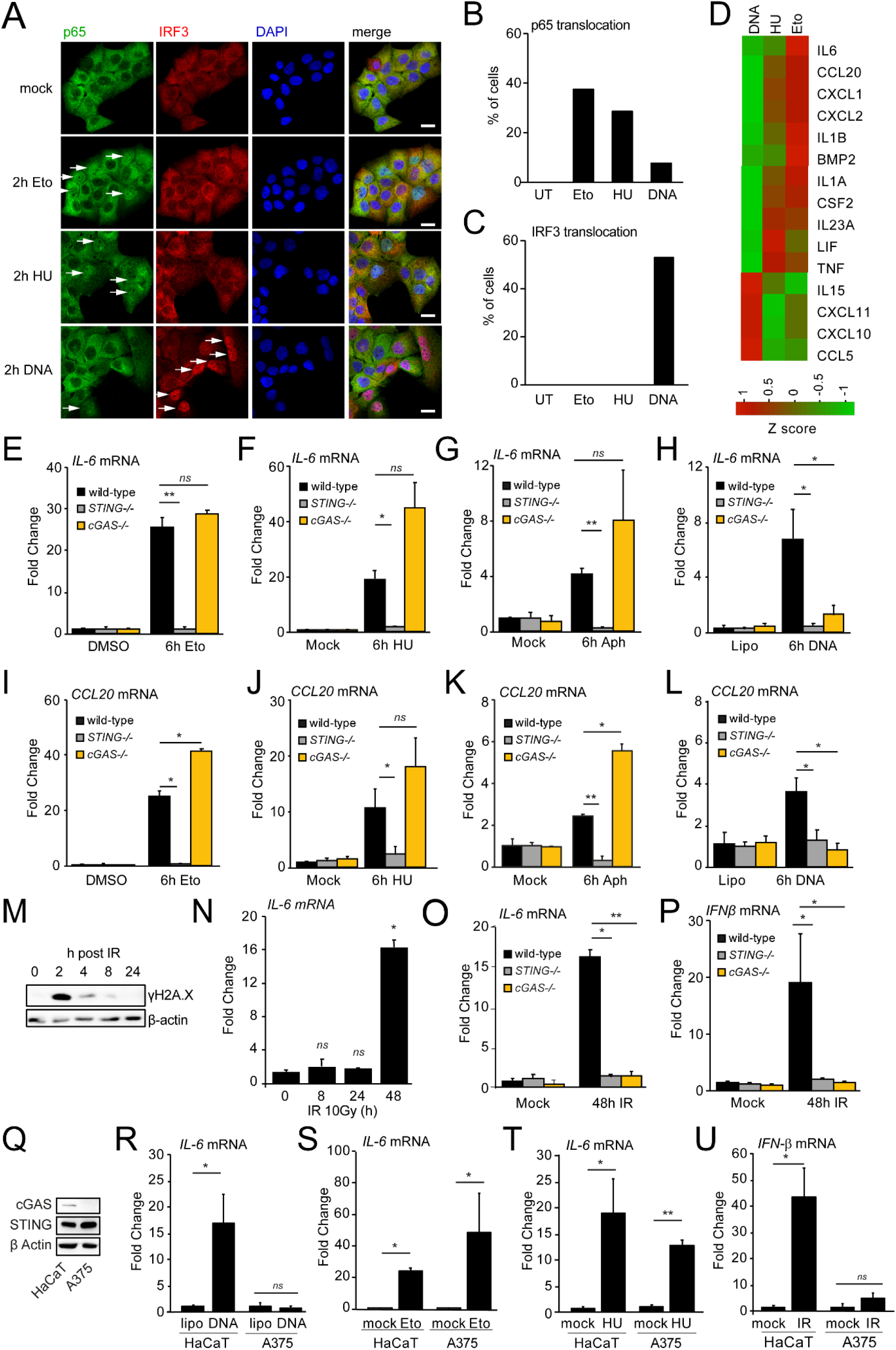
Replication stress induces a cGAS-independent, but STING-dependent inflammatory response. A: HaCaT keratinocytes were grown on coverslips and treated with 50 μM etoposide (Eto), 5mM hydroxyurea (HU) or transfected with 5μg/ml herring testis (HT) DNA for 2h. Cells were fixed and stained for endogenous NF-κB p65 (green), IRF3 (red) and DNA (DAPI, blue). Scale bar: 20μm. B, C: Cells stained for IRF3 and p65 from experiments as in A were scored for predominant nuclear localisation. At least 100 cells were scored per experiment. D: qPCR array for the quantification of cytokine and chemokine mRNA expression in HaCaT cells treated with 1 μg/ml transfected DNA, 5mM HU or 50μM etoposide for 6h. Fold induction was compared to untreated cells. E-L: qPCR analysis of IL-6 and CCL20 mRNA expression in cGAS or STING-deficient HaCaT cells or wild type controls, treated for 6h with DMSO or 50μM etoposide (E, I), 5mM HU (F, J), or 10μM aphidicolin (G, K). As control, cells were mock transfected (Lipo) or transfected with 1μg/ml herring testis DNA (H, L). qPCR experiments were carried out on triplicate samples, and IL-6 or CCL20 mRNA values were normalised to GAPDH mRNA levels and expressed as fold changes compared to mock treated wild type controls. M: HaCaT cells were exposed to ionising radiation (IR, 10 Gy), and the phosphorylation of histone γH2A.X was monitored by immunoblotting following recovery for up to 24h as indicated. N: HaCaT cells were exposed to IR (10Gy) and following recovery for the times indicated, and the expression of IL-6 mRNA was quantified by qRT-PCR. P, O: qRT-PCR analysis of IL-6 or interferon-β (IFN-β) mRNA from cGAS- or STING-deficient HaCaT cells and wild type controls, 48h post recovery from IR exposure (10Gy). Q: Immunoblotting for STING and cGAS protein expression in HaCaT cells or A375 melanoma cells, with β-actin control. R-T: HaCaT cells or A375 cells were transfected with 1μg/ml herring testing DNA, treated with 50μM etopside (Eto) or 5mM hydroxyurea (HU) for 6h, and the expression of IL-6 mRNA was quantified by qRT-PCR, normalised to GAPDH mRNA and expressed as fold change over respective mock treatments. U: HaCaT or A375 cells were exposed to ionizing radiation (IR, 10Gy), and IFN-β mRNA expression was quantified by qRT-PCR after recovery for 48h. Data from qRT-PCR experiments are shown as mean values of triplicate samples; error bars represent SD. Experiments were performed at least three times. Statistical significance was calculated using unpaired Student’s t-tests; * p<0.05, ** p<0.01, *** p<0.001.

We found that the cytokine/chemokine response to HU and etoposide occurred within 6 h of treatment, with HU causing a further increase in IL-6, CCL20 and IFN-β mRNA expression by 24h (Suppl. Fig. 1A-D). Using cGAS or STING KO HaCaT keratinocytes generated by CRISPR-Cas9-based gene targeting^32^, we find that the expression of IL-6 and CCL20 mRNA in response to etoposide or HU treatment, or treatment with the DNA polymerase inhibitor aphidicolin is dependent on STING, but not on cGAS (Fig. 1E-G and I-K). In contrast, DNA transfection-driven induction of IL-6 and CCL20 was dependent on both STING and cGAS, as expected for the cytosolic DNA sensing response (Fig. 1H and L).^12^

As replication fork stalling can ultimately lead to the formation of DSBs, we tested whether direct DSB formation caused by ionizing radiation (IR) also induces an acute inflammatory response. We found that while exposure to 10Gy IR caused the expected increase in the DNA damage marker γH2A.X (Fig. 1M), we did not detect induction of IL-6 or interferon-β (IFNβ) mRNA until 48h post recovery from IR (Fig. 1N, Suppl. Fig. 1E). This delayed response to IR-induced DNA damage was dependent on both cGAS and STING (Fig. 1O and P). Delayed cGAS-STING activation has previously been attributed to the detection of cytosolic DNA from post-mitotic micronuclei which form several days after recovery from IR-induced damage^9,10^, although this has been disputed^33^. Our data demonstrates that there are distinct responses to genotoxic agents that drive cytokine induction: a delayed cGAS-dependent response to IR-induced DNA damage, and an earlier cGAS-independent response to replication stress-inducing agents such as HU, aphidicolin and etoposide.

The cell-intrinsic innate immune response to DNA damage and replication stress may play a role in cancer cells which frequently display elevated levels of genotoxic stress, increased further by treatment with replication stress-inducing chemotherapies^34^. This in turn has the capacity to modulate the tumour micro-environment for pro- and anti-tumour immune functions^35^. We found that the inflammatory response to replication stress was preserved in several cancer cell lines, including in the human bladder cancer cells T24 and J82 which secreted IL-6 in response to HU and etoposide treatment (Suppl. Fig. 1F and G). As observed in HaCaT cells, treatment with HU or etoposide caused the nuclear translocation of NF-κB p65 in T24 bladder cancer cells, while DNA transfection resulted in the predominant activation of IRF3 (Suppl. Fig. 2A and B). Pre-treatment of T24 cells with cGAS siRNA or the cGAS inhibitor C-76 inhibited the activation of IRF3 following DNA transfection, but did not affect the response to etoposide or HU (Suppl. Fig. 2A and B). An inflammatory response to replication stress was also observed in A375 melanoma cells. These cells do not express cGAS, and therefore are unable to respond to transfected dsDNA, as described previously^36^ (Fig. 1Q and R). However, IL-6 mRNA was robustly induced in A375 melanoma cells following treatment with etoposide or HU (Fig. 1S and T), providing further confirmation that the acute inflammatory response to replication stress is cGAS-independent. In line with our data from cGAS-deficient HaCaT cells, A375 cells were also unable to induce a delayed IFN-β response 48h post IR treatment (Fig. 1U), providing additional evidence that the responses to IR-induced DNA damage and replication stress are distinct.

### IFI16 is required for the proinflammatory cytokine response to replication stress

Since replication stress-driven cytokine induction is cGAS-independent, we hypothesised that the DNA binding protein IFI16 may act as a replication stress sensor in this context. In human cells, IFI16 is a predominantly nuclear protein which shuttles between the nucleus and cytosol and interacts with STING to promotes its activation in response to both cytosolic DNA and to etoposide-induced damage27,31. To test the involvement of IFI16 in replication stress-driven inflammation, we use IFI16 KO HaCaT cells^32^ to assess IL-6 mRNA levels by qRT-PCR and IL-6 secretion by ELISA. Treatment of WT HaCaT cells with HU or etoposide resulted in increased IL-6 mRNA levels by 6h, and increased IL-6 cytokine secretion by 12h post HU treatment (Fig. 2A-D). IL-6 mRNA expression was impaired in cells lacking either IFI16 or STING (Fig. 1A-C). IFI16-deficient cells were also unable to secrete IL-6 cytokine at 12h and 24h after treatment with HU (Fig. 2D).

**Figure 2:**
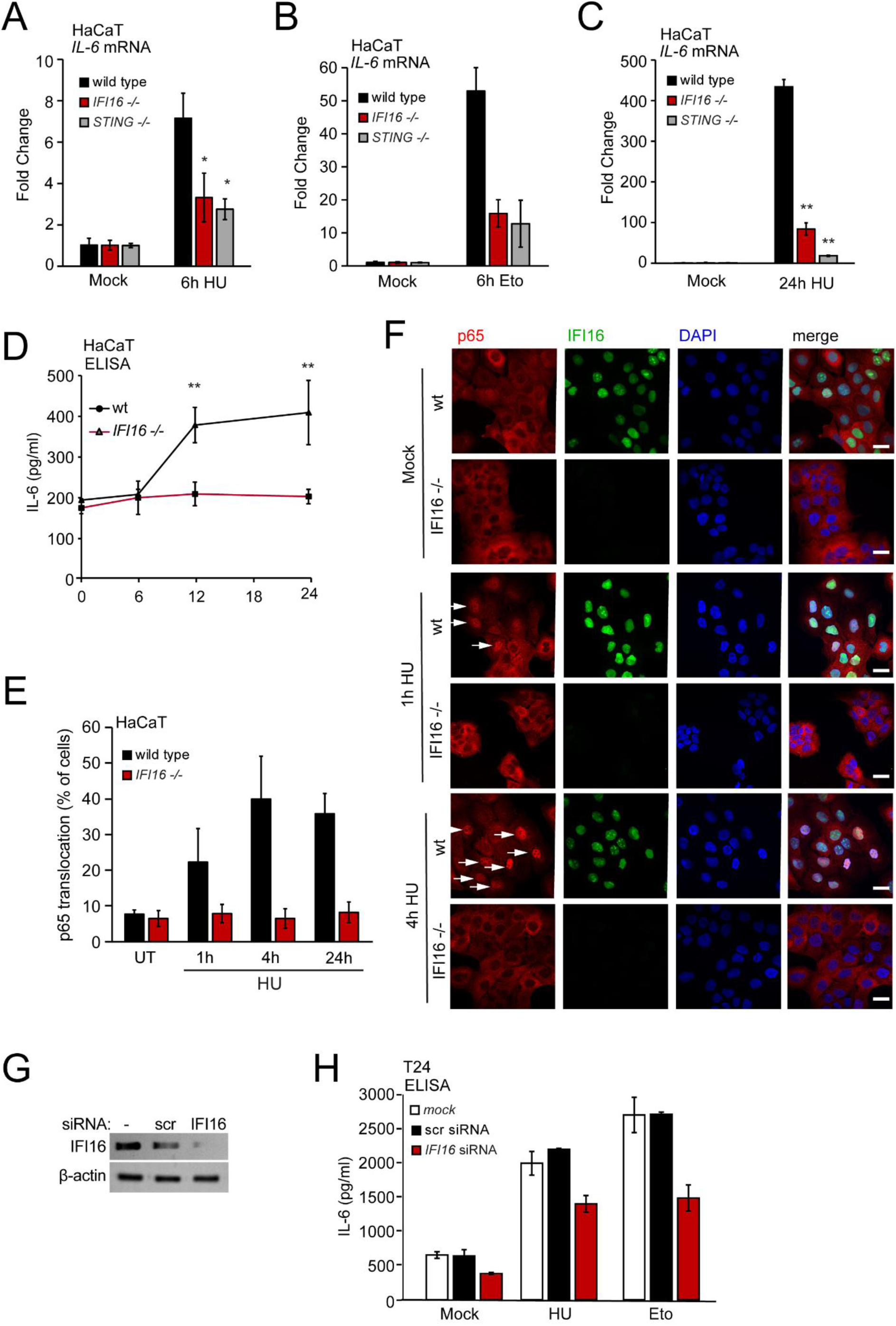
IFI16 is required for the innate immune response to replication stress. A, B: WT, IFI16−/− or STING−/− HaCaT cells were treated with 5mM HU (A) or 10, 50 μM etoposide for 6h. IL-6 mRNA expression was quantified by qRT-PCR, normalised to GAPDH mRNA and expressed as fold induction over mock-treated control cells. C: WT, IFI16−/− or STING−/− HaCaT cells were treated with 5mM HU for 24h, and IL-6 mRNA expression was quantified by qRT-PCR as in A. D WT and IFI16−/− HaCaT cells were treated with 5mM HU for the times indicated, and the secretion of IL-6 protein in cell supernatants was quantified by ELISA, based on a standard curve of recombinant IL-6. E-F: WT or IFI16−/− HaCaT cells were grown on coverslips, mock treated or treated with 5mM HU for the times indicated. Cells were fixed and stained for endogenous NF-κB p65 (red), IFI16 (green) and DNA (DAPI, blue) and imaged by confocal microscopy. Scale bar: 20μm.The percentage of cells displaying predominantly nuclear NF-κB p65 in E, scoring a minimum of 100 cells per sample. G: T24 bladder cancer cells were transfected with non-targeting control siRNA, or an siRNA targeting IFI16. After 48h, cells were treated with etoposide (Eto, 50μM) or hydroxyurea (HU, 5mM) for 24h. Secreted IL-6 protein was quantified by ELISA.

We also monitored the nuclear translocation of the NF-κB p65 subunit following HU-induced replication stress by immunofluorescence/confocal microscopy. In wild type HaCaT cells, p65 nuclear translocation was detectable from 1h post HU treatment, with approximately 30-40% of cells displaying nuclear p65 staining by 4h, and this was sustained to 24h post treatment (Fig. 2E and F). p65 nuclear translocation was impaired in IFI16-deficient cells at all timepoints tested (Fig. 2E and F). We also depleted IFI16 in T24 bladder cancer cells using RNAi (Fig. 2G). IFI16 depletion reduced the amount of IL-6 cytokine secreted in response to etoposide or HU treatment of these cells, when compared with cells that were mock transfected or transfected with scrambled control siRNA (Fig. 2H). Thus, we conclude that IFI16 is required for the early inflammatory response to replication stress.

### Replication stress is sensed prior to DSB formation

IFI16 is a DNA binding protein which interacts with nucleosome-free linear dsDNA and has also been reported to bind to supercoiled and cruciform DNA, and structured DNA ligands^19,30–32^. Thus, we investigated which DNA features might be detected by IFI16 as damage- or stress-associated molecular patterns (DAMPs or SAMPs) during replication stress. Treatment with replication stress inducers induces the remodelling of replication forks (fork reversal) but can also cause fork collapse and DNA breaks^37^. Given that we observe IFI16-dependent p65 activation within the first hours of HU treatment (Fig. 2G and H), cytokine mRNA expression from 6h, and IL-6 secretion from 12h post-treatment in HaCaT cells (Fig. 2A-F), we tested whether these endpoints are preceded by the generation of DSBs. Using Comet assays, we found that HU treatment did not cause a significant increase in DSBs within the 6h period post-treatment, although increased DSB formation was apparent after 24h (Fig. 3A). However, even at this time point, the extent of DSB formation was much lower than that induced by a 6 or 24h treatment with etoposide (Fig. 3A). We also did not observe any increase in micronuclei formation within 24h of HU treatment (Fig. 3B). To further examine to what extent the inflammatory response is linked to the accumulation of DNA damage in individual HU-treated cells, we monitored the timing of p65 translocation in T24 bladder cancer cells, in parallel with quantification of γH2A.X phosphorylation as a DNA damage marker (Fig. 3C). γH2A.X phosphorylation was readily detectable after 24h HU treatment, but in the first 6h post-HU treatment γH2A.X phosphorylation did not exceed basal levels in the vast majority of cells (Fig. 3C and D). As we observed in HaCaT cells, the timing of NF-κB p65 translocation is not chronologically linked to increased levels of DNA damage. Instead, we observe an early wave of p65 nuclear translocation, starting at 1-2h post HU treatment and detectable in approximately 30% of cells between 2h and 6h post HU treatment, with no further increase at later time points when DNA damage is more prominent (Fig. 3C-E). Overall, our data suggest that the innate immune response to replication stress is initiated prior to DNA DSB formation, and thus likely involves the recognition of replication stress, rather than overt DNA damage.

**Figure 3:**
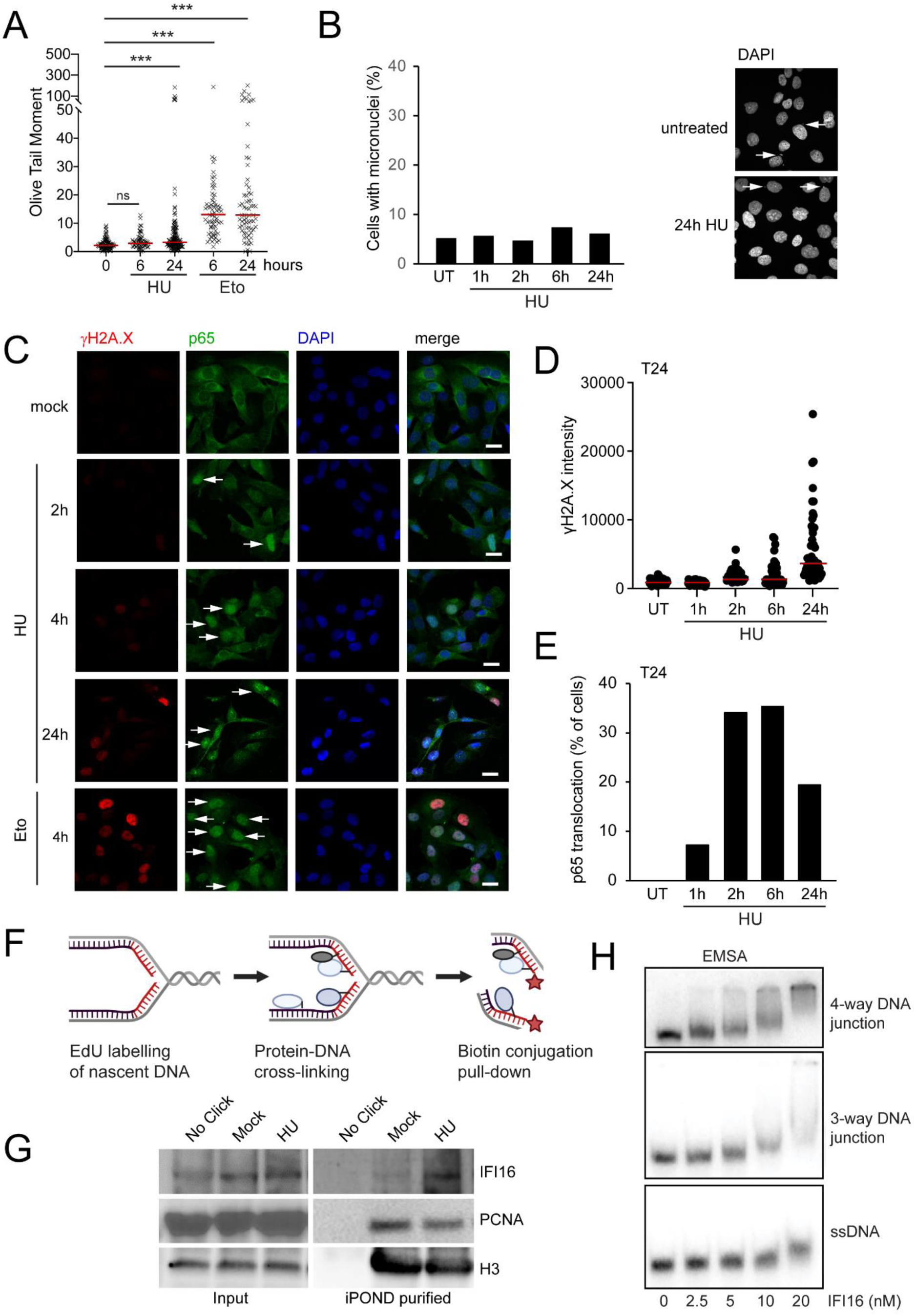
IFI16 senses stalled replication forks prior to the generation of replication stress-induced DNA damage. A: Comet assay for the quantification of double-strand DNA breaks in HaCaT cells treated with 5mM HU or 50μM Eto for the times indicated. Shown are calculated Olive Tail Moment values for individual nuclei; bars represent median values. B: HaCaT cells were treated with 5mM HU for the times indicated, fixed and stained with DAPI. The presence of micronuclei in the cytosol was scored by confocal microscopy analysis of >100 cells per sample. C-E: T24 bladder cancer cells were treated with 5mM HU or 50μM Eto for the times indicated, fixed and stained for phosphorylated γH2A.X (red), NF-κB p65 (green) and DNA (DAPI, blue). Scale bar: 20μm. D: Nuclear γH2A.X staining intensity from cells treated with 5mM HU for the times indicated and stained as in (C), quantified as mean values in individual nuclei, with bars representing median values of at least 50 cells per sample. E: Cells treated with 5mM HU as in C were scored for nuclear translocation of endogenous NF-κB p65. P65 translocation is expressed as the percentage of cells displaying predominantly nuclear p65 localisation in the same samples as D. F: Schematic of the iPOND approach for the identification of proteins that associate with nascent DNA at replication forks. G: U2OS cells were mock-treated or treated with 3 mM HU. After 2 hrs, cells were labelled with EdU for 15 min, and the iPOND procedure was performed. A ‘no click’ reaction was included to as a control. iPOND eluates were resolved by western blotting and probed with the indicated antibodies. H: Recombinant IFI16 was incubated with single-stranded (ss) DNA, a 3-way DNA junction or 4-way DNA junction as indicated. IFI16 binding to each target was assessed by EMSA with recombinant IFI16. All experiments were performed at least 3 times. Where indicated, statistical significance was calculated using unpaired Student’s T tests; * p<0.05, ** p<0.01, *** p<0.001.

### IFI16 binds to stalled replication forks

Given that IFI16 is a DNA binding protein which is located predominantly in the nucleus, we hypothesised that IFI16 might bind directly to stalled replication forks. Therefore, we first tested whether IFI16 directly associates with stalled replication forks during replication stress, by employing the Isolation of Proteins on Nascent DNA (iPOND) technique^26^ to purify protein complexes associated with EdU-labelled nascent DNA (Fig. 3F). We performed the iPOND technique in HaCaT cells, which were either left untreated or were treated with HU for 2h. We observed association between IFI16 and nascent DNA in untreated cells, and this association was elevated upon HU-mediated induction of replication stress (Fig. 3G). Thus, our data demonstrates the physical association of IFI16 and nascent DNA at stalled replication forks.

To investigate whether IFI16 can bind fork structures directly, we generated recombinant IFI16 in the *Sf*9 baculoviral system (Suppl. Fig. 2A) and conducted electrophoretic mobility shift assays (EMSA) to assess the relative *in vitro* binding of IFI16 to ssDNA, 3-way dsDNA junctions (which mimic replication forks) and 4-way dsDNA junctions (which mimic the Holliday Junctions formed by fork reversal). IFI16 bound poorly to ssDNA but exhibited high-affinity binding to a dsDNA oligo as expected (not shown), and to both 3-way and 4-way dsDNA junctions (Fig. 3H).

### IFI16 protects reversed forks from DNA2 and MRE11-mediated degradation

Given that is recruited to stalled replication forks and directly binds DNA in vitro, we hypothesised that IFI16 has additional functions at the replication fork DNA, independent of its role in innate immune activation. To test whether IFI16 affects the cell’s ability to protect or repair replication stress-induced DNA breaks, we monitored γH2A.X phosphorylation following HU-induced replication stress in wild type and IFI16-deficient HaCaT cells. Pre-extraction was used to remove soluble or loosely bound nuclear proteins. We found that in the absence of IFI16, cells displayed markedly higher levels of DNA damage as indicated by γH2A.X phosphorylation following HU treatment (Fig. 4A and B). This effect could be observed as early as 4h post HU treatment and persisted at 24h, showing that IFI16 protects stalled replication forks from replication stress-induced damage.

**Figure 4:**
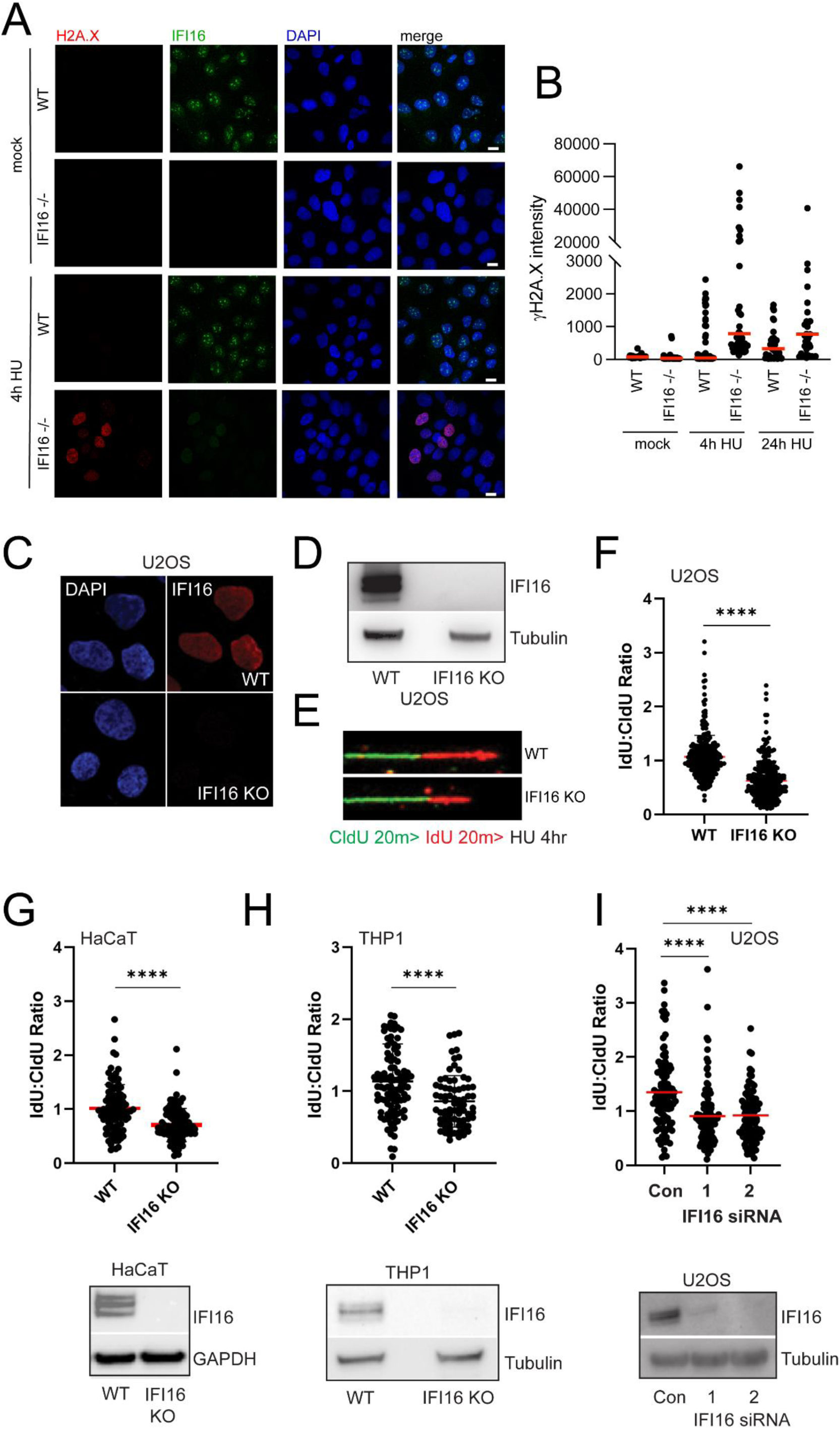
Loss of IFI16 leads to nascent DNA degradation during replication stress. A-B: WT or IFI16 −/− HaCaT cells were grown on coverslips and treated with 5mM HU for the times indicated. Cells were pre-extracted, fixed, and stained for phosphorylated γH2A.X (red), IFI16 (green) or DNA (DAPI, blue). Nuclear γH2A.X staining intensity was quantified in individual nuclei, with bars representing median values of at least 50 cells per sample. C-D: IFI16 KO U2OS cells were generated via lentiviral delivery of Cas9 and IFI16 gRNA. Confirmation of IFI16 loss was confirmed via both immunofluorescence (C) and Western blotting (D). E-F: DNA fibre assay to assess fork protection. WT and IFI16 KO U2OS cells were sequentially labelled with CldU and IdU as indicated, and then treated with 4 mM HU for 4 hrs. Cells were then lysed, and DNA fibres spread, fixed and stained, then imaged by confocal microscopy. CldU and IdU tract lengths were measured using ImageJ, and the IdU:CldU ratio of each fibre determined. Reduced IdU:CldU ratio is indicative of nascent DNA degradation. G-I: WT and IFI16 KO THP1 monocytes, IFI16-depleted U2OS cells, or WT and IFI16 KO HaCaT cells were treated and analysed as in (E). All experiments were performed three times. Statistical significance was calculated using unpaired t-tests or one-way ANOVA: *** p<0.001, **** p<0.0001.

To test whether the association of IFI16 protects stalled replication forks from degradation, we employed a fork protection assay, in which cells are sequentially labelled with the halogenated nucleoside analogues CldU and IdU, followed by chronic fork stalling in the presence of high-dose HU. The lengths of contiguous CldU and IdU tracts were then measured and expressed as a ratio. Under these conditions, any reduction in IdU:CldU ratio can be considered a consequence of nucleolytic degradation of nascent DNA. We assessed fork protection in several different models of IFI16 loss-of-function. Firstly, we generated IFI16 KO U2OS cells via CRISPR-Cas9-mediated gene targeting and employed existing IFI16 KO THP1 monocytes and IFI16 KO HaCaT keratinocytes, as well as U2OS cells depleted of IFI16 via RNAi. IFI16 loss or depletion was assessed by Western blotting and/or immunofluorescence (Fig. 4C-I). We observed significant reductions in IdU:CldU ratio in all IFI16 KO backgrounds relative to paired WT cells, as well as in IFI16-depleted U2OS cells relative to cells transfected with a non-targeting control siRNA (Fig. 4C-I). This provides robust evidence that IFI16 binding protects stalled replication forks from degradation.

To test whether fork reversal is a pre-requisite for the loss of nascent DNA integrity in HU-treated IFI16 KO cells, we performed the fork protection assay in WT and IFI16 KO U2OS cells depleted of the fork reversal factors SMARCAL1 and ZRANB3. Depletion of both factors was confirmed by Western blotting (Suppl. Fig. 2B). As before, loss of IFI16 resulted in a significant reduction in IdU:CldU ratio in cells treated with a non-targeting control siRNA, but this phenotype was rescued by depletion of either SMARCAL1 or ZRANB3 (Fig. 5A). This indicates that nascent DNA degradation in IFI16 KO cells lies downstream of fork reversal^38,39^.

**Figure 5:**
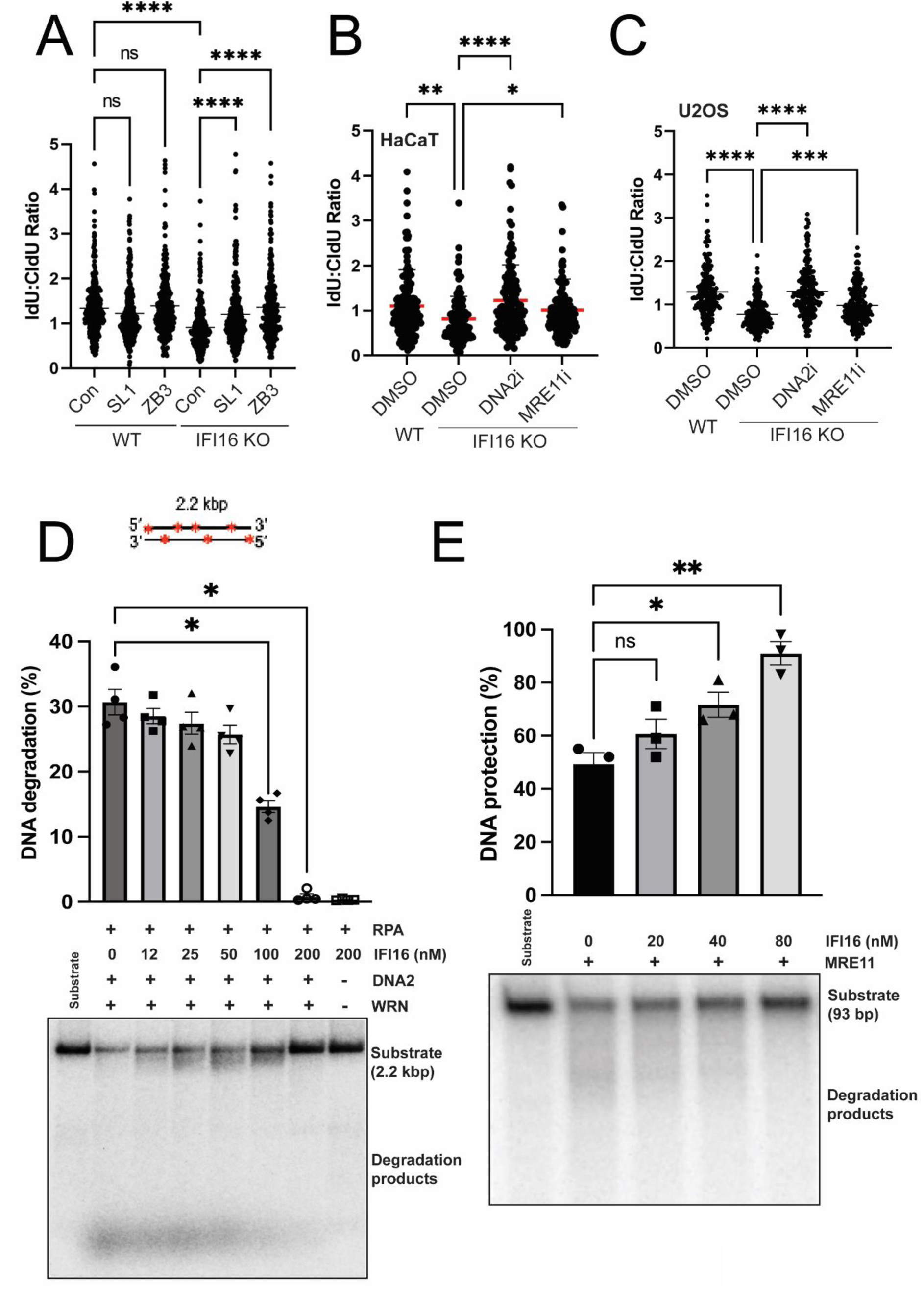
IFI16 protects reversed forks from DNA2 and MRE11-mediated degradation. A: WT and IFI16 KO U2OS cells were transfected with a control siRNA, or an siRNA targeting either SMARCAL1 or ZRANB3. Cells were then treated, assessed by DNA fibre assay and data analysed as in Fig 3B. B and C: WT and IFI16 KO U2OS were pre-treated with either DNA2 inhibitor C5 (20 μM) or the MRE11 exonuclease inhibitor Mirin (25 μM) and assessed as in (A) following dual-labelling and incubation with 4 mM HU for 4 hrs. D: DNA end resection assay with DNA2 (30 nM), WRN (30 nM) and RPA (176 nM) in the presence of various concentrations of recombinant IFI16 (12, 25, 50, 100, 200 nM) on a randomly labelled 2.2-kbp dsDNA. Top, schematic of the 2.2-kbp dsDNA substrate. Middle, quantitation of DNA degradation. Averages shown, n = 4; error bars, SEM. Bottom, representative gel from four independent experiments. E: Exonuclease assay with recombinant MRE11 (400 nM) and various concentrations of recombinant IFI16 (0, 20, 40 and 80 nM) on a 5′ end-labelled 93-bp dsDNA. Top, quantitation of DNA protection. Averages shown, n = 3; error bars, SEM. Bottom, representative gel from three independent experiments. All experiments were performed three times. Statistical significance was calculated using unpaired t-tests or one-way ANOVA: ns=not significant, * p<0.05, **** p<0.0001.

To assess nuclease involvement in nascent DNA degradation in IFI16 KO cells, we performed the fork protection assay in IFI16 KO U2OS or HaCaT cells following pre-treatment with either the MRE11 exonuclease inhibitor mirin or the DNA2 inhibitor C5. Inhibition of either DNA2 or MRE11 resulted in rescue of fork protection in IFI16 KO cells of both backgrounds, suggesting direct involvement of these nucleases in the degradative process antagonised by IFI16 (Fig. 5B and C). To test whether IFI16 can directly repress nuclease activity *in vitro*, we performed MRE11 exonuclease digestion assays on a dsDNA substrate in the presence of increasing concentrations of recombinant IFI16. We also tested the ability of IFI16 to modulate the WRN-assisted digestion of a dsDNA substrate by DNA2. Incubation with nanomolar levels of recombinant IFI16 resulted in inhibition of dsDNA digestion by either WRN-DNA2 and MRE11 *in vitro* (Fig. 5D and E).

### IFI16 is required for IFNβ-driven rescue of fork protection in BRCA-deficient cells

The canonical tumour suppressors BRCA1 and BRCA2 protect reversed forks^7^. Recent reports demonstrate that IFN-β supplementation rescues fork protection in BRCA-deficient cell models^14^. This constitutes a link between innate immunity and fork protection, as IFN-β is produced as a response to viral infections and DNA damage, and this in turn up-regulates several innate immune sensor proteins including IFI16^17,40^.

Since IFI16 binds and protects DNA at reversed forks and is potently induced by type I IFNs in many cell types^17^, we hypothesised that IFI16 may contribute to IFN-mediated fork protection in BRCA-deficient cells. To test this, we assessed fork protection in WT and IFI16 KO U2OS cells following siRNA-mediated depletion of BRCA1 or BRCA2, in the presence and absence of IFN-β supplementation. IFNβ treatment led to robust induction of IFI16 in these cells (Fig. 6A), and the depletion of BRCA1 and BRCA2 in IFN-β-treated cells was confirmed by Western blotting (Fig. 6B). As previously reported, BRCA1 or BRCA2 depletion resulted in nascent DNA degradation in WT U2OS cells, as did loss of IFI16. In agreement with the findings of the Penengo laboratory, IFN-β supplementation completely rescued defective fork protection in BRCA1 or BRCA2-depleted WT U2OS cells (Fig. 6C)^14^. However, this effect was not observed in IFI16 KO cells (Fig. 6C), demonstrating that IFN-mediated restoration of fork protection is IFI16-dependent. Furthermore, we did not observe an additional increase in DNA degradation in BRCA1-depleted IFI16 KO cells (Fig. 6C), suggesting that BRCA1 and IFI16 may function in the same pathway. We did observe a significant decrease in nascent DNA degradation in BRCA2-depleted IFI16 KO cells relative to controls (Fig. 6C), suggestive of a mechanistic difference between BRCA1 and BRCA2-mediated fork protection. We next hypothesised that IFI16 overexpression can circumvent loss of fork protection in BRCA1 or BRCA2-deficient cells, even in the absence of IFNs. To test this, we depleted BRCA1 or BRCA2 in HEK293 cells, or in HEK293 cells stably expressing IFI16-FLAG. BRCA1/2 depletion and IFI16-FLAG expression were confirmed by Western blotting (Fig. 6D). Depletion of BRCA1 or BRCA2 led to a significant increase in nascent DNA degradation in HEK293 cells, which was completely abolished in cells over-expressing IFI16-FLAG (Fig. 6E). Overall, this data indicates that IFI16 is an important component of the IFN-mediated restoration of fork protection in BRCA-deficient cells, lending further support to our observation that IFI16 prevents the degradation of nascent DNA during replication stress.

**Figure 6:**
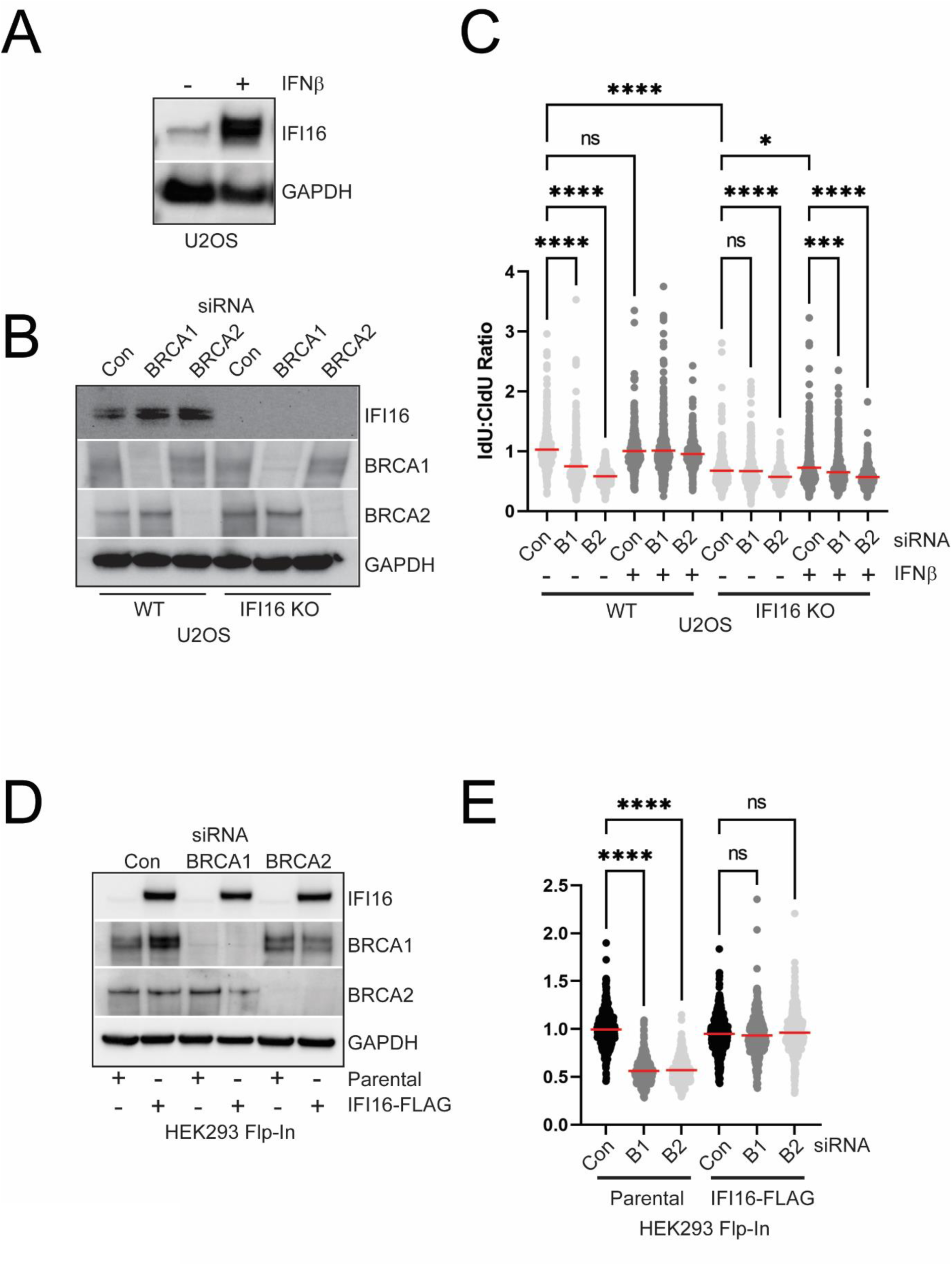
IFI16 is required for IFNβ-driven rescue of fork protection in BRCA-deficient cells. A: WT U2OS cells were treated with 30 U/ml recombinant IFN-β. After 48 hrs, cells were lysed, and IFI16 levels assessed by Western blotting with the indicated antibodies. B and C: WT or IFI16 KO U2OS cells were transfected with a control siRNA, or siRNAs targeting either BRCA1 or BRCA2, and treated with 30 U/ml recombinant IFN-β. After 48 hrs, cells were sequentially CldU and IdU-labelled and treated with 4 mM HU for 4 hrs. As previously, IdU:CldU ratios were calculated, and the data displayed in (B). BRCA1 and BRCA2 depletion and IFI16 loss were validated by Western blotting (C). D and E: Parental Flp-In HEK293 cells, or HEK293 cells stably expressing IFI16-FLAG were transfected with a control siRNA, or siRNAs targeting either BRCA1 or BRCA2. IFI16 expression and BRCA1/2 depletion were confirmed by Western blotting (D). After 48 hrs, cells were sequentially CldU and IdU-labelled and treated with 4 mM HU for 4 hrs. As previously, IdU:CldU ratios were calculated, and the data displayed in (E). All experiments were performed three times. Statistical significance was calculated using one-way ANOVA: ns=not significant, * p<0.05, ***p<0.001 and **** p<0.0001.

Overall, our findings demonstrate that IFI16 binds directly to nascent DNA at stalled replication forks where it acts both as fork protection factor, and as an innate immune sensor for replication stress. In this way, IFI16 provides a further link between innate immunity and genomic stability.

## Discussion

DNA damage and genome instability can initiate a cell-intrinsic inflammatory response in both untransformed and cancerous cells. However, the molecular features that are detected as damage- or stress-associated molecular patterns under these circumstances remain to be defined. A delayed response several days after recovery from DNA damage has been attributed to the detection of cytosolic DNA from micronuclei by the DNA sensor cGAS^9,10^, but recent data have questioned this notion^33^. Here, we demonstrate that stalled replication forks are detected as a danger signal by the innate immune system and initiate an NF-κB-mediated inflammatory response within the first hours following replication stress. We show that the detection of replication stress is cGAS-independent, but STING-dependent, and mediated by the DNA binding protein IFI16. We find that IFI16 directly binds to nascent DNA at stalled replication forks and initiates an innate immune response prior to the formation of replication stress-induced DNA damage. Thus, we propose that stalled forks are detected as stress-associated molecular patterns (SAMPs), with IFI16 and STING acting as guard proteins to alert the cell to a disruption of normal cellular processes. The notion that innate immune guard proteins monitor cellular functions for signs of disruption by pathogens (so-called effector-triggered immunity^41^) has recently been proposed, and may have applications beyond infection. For instance, the detection of transcription stress and ribotoxic stress due to translation block have recently been shown to contribute to the inflammatory response to UV irradiation and other cellular insults^42,43^. Our work has now added DNA replication stress as another transient disruption of cellular processes which acts as innate immune alarm signal prior to cell damage. The detection of stalled replication forks may of relevance during neoplastic transformation and oncogene-induced replication stress, which are known drivers of inflammation. It remains to be tested whether stalled replication forks can also be detected as SAMP during viral infection, for instance under conditions where the availability of DNA replication components becomes limiting.

In addition to the innate immune function of IFI16 in the detection of reversed replication forks, we also describe a novel function for IFI16 in the protection of reversed fork DNA from nucleolytic degradation. This finding mirrors prior observations that loss of the DNA sensor cGAS leads to nascent DNA degradation during replication stress and altered fork progression^44^. Protection of reversed replication forks from nuclease-mediated degradation is an important component of the cellular response to replicative stress. To date, a number of replication fork protection factors have been identified, including the canonical tumour suppressors BRCA1 and BRCA2, and the nuclease interactor MRNIP^6,7,38,45^. Here, we identify the innate immune DNA sensor IFI16 as an additional fork protection factor, which binds directly to nascent DNA at stalled, reversed replication forks and limits degradation by the nucleases DNA2 and MRE11.

Recent work from the Penengo laboratory demonstrates that IFN supplementation fully rescues nascent DNA degradation associated with loss of function of BRCA1 or BRCA2^14^. This rescue is dependent on the ubiquitin-like modifier Interferon-Stimulated Gene 15 (ISG15), which has been independently linked to the mitigation of replication stress^46^. We demonstrate that IFN-mediated restoration of fork protection in BRCA1 or BRCA2-depleted cells is dependent on IFI16. At present, the mechanism via which IFN/ISG15-mediated restoration of fork protection occurs is unclear. Type 1 IFNs induce the transcription of hundreds of interferon-stimulated genes^40^, although we note that only a handful of these factors have also been identified in iPOND eluates^26^. We hypothesise that the direct association of IFI16 with nascent DNA following IFN treatment can facilitate compensatory protection from nucleolytic activities in the absence of functional BRCA1/2-dependent RAD51 loading. It is possible that RAD51 limits IFI16 association with nascent DNA. RAD51 binds both dsDNA and ssDNA, and BRCA2-driven dsDNA binding by RAD51 has recently been demonstrated to drive fork protection^47^. IFI16 binds more avidly to dsDNA, and therefore any competition between RAD51 and IFI16 is more likely to occur in this context. Work from the Petrini laboratory shows that the inducible loss of murine *Nbs1* leads to enhanced association of the IFI16 homologues *Ifi204* and *Ifi205* with nascent DNA, further suggesting that IFI16 responds dynamically to replication stress^46^. It is also possible that global chromatin alterations underpin the effect of IFNs on replication fork protection, since IFN treatment has been extensively linked with alterations to chromatin architecture^48^ and IFI16 modulates chromatinization of viral DNA genomes^49,50^. Our work also demonstrates that even in the absence of IFNs, IFI16 overexpression rescues nascent DNA degradation in BRCA1 or BRCA2-depleted cells. This implies that increased levels of IFI16 alone are capable of restoration of fork protection, and that this process occurs independently of the complex events that occur downstream of IFN signalling. Given that the IFI16 gene locus is frequently amplified in cancers, it is conceivable that IFI16 upregulation acts as a compensatory mechanism for some tumours, allowing them to cope with elevated levels of replication stress. Overall, our study highlights the complex interplay between replication stress, DNA damage and innate immunity, with IFI16 acting as a mediator in the regulation of both the inflammatory response to replication stress, and the stability of stalled replication forks.

## Data and materials availability

All data needed to evaluate the conclusions in the paper are present in the paper. Additional data related to this paper may be requested from the authors.

## Acknowledgements

We thank Dr Andrew Pierce, Dr Panagiotis Kotsantis and Dr Edgar Hartsuiker for useful discussions.

## Author contributions

L.U., C.J.S. and T.A.W. conceived the study. A.G., T.A.W., O.P.G.W., L.G.B., C.M.J., E.G.V., V.T., J.P.M., J.R. and T.D.J.W. designed and performed experiments and analysed data. S.H. prepared recombinant IFL16. I.C. and D.B. performed the *in vitro* experiments supervised by P.C. L.U. supervised the innate immunity aspects of this work and C.J.S. supervised the work relating to the functions of IFI16 at the replication fork. L.U. and C.J.S. obtained funding and wrote the manuscript with input from all the authors.

---

**Supplementary Figure 1.**
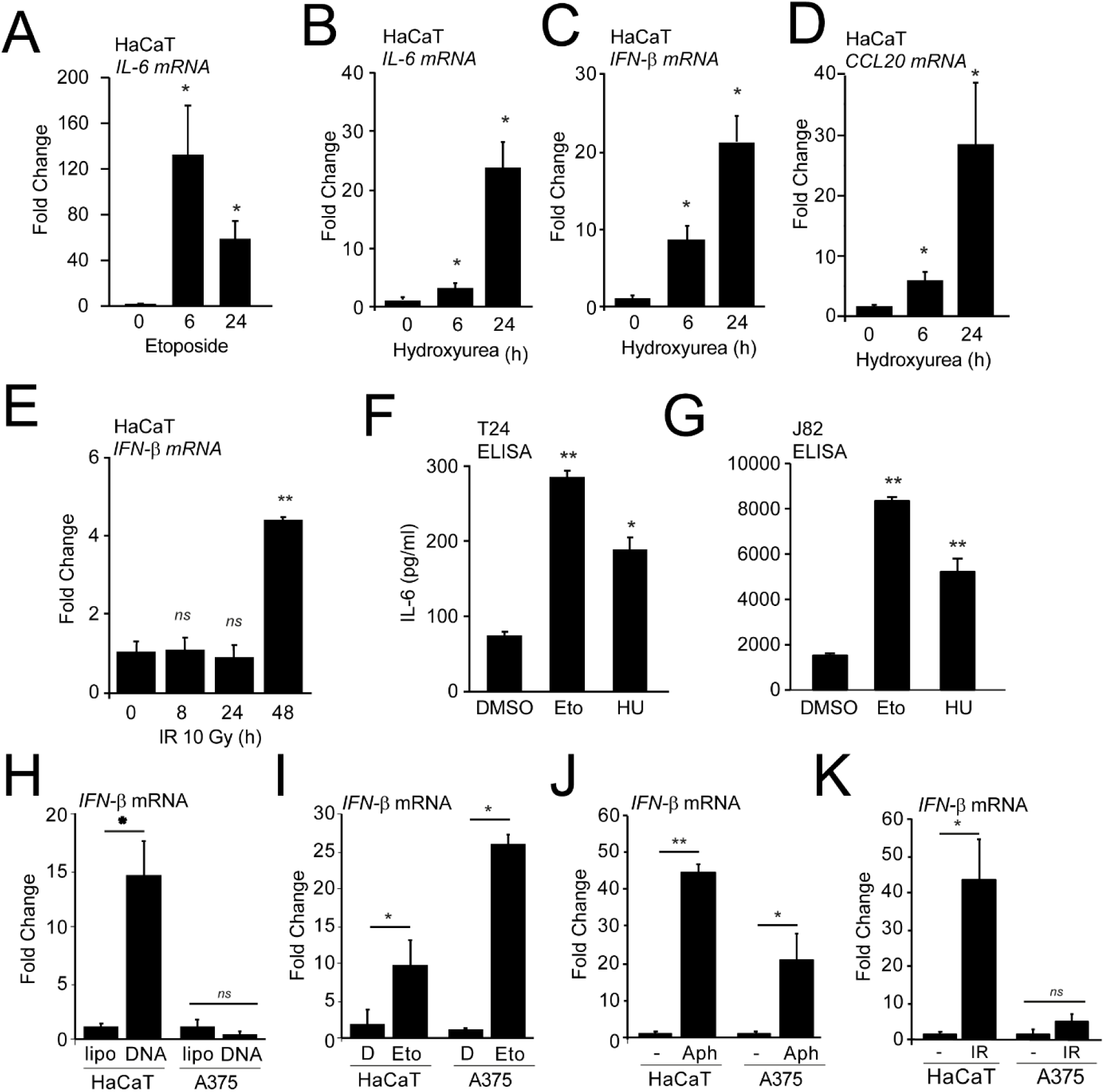
A-D: HaCaT keratinocytes were treated with 50 μM etoposide (Eto) or 5mM hydroxyurea (HU) for the times indicated, and the expression of IL-6 (A, B), IFN-β (C) or CCL20 mRNA (D) was analysed by qRT-PCR, normalised to β-actin mRNA and expressed as fold induction over mock treated cells. E: HaCaT cells were treated with ionizing radiation (IR, 10 Gy) and IFN-β mRNA was quantified by qRT-PCR at the indicated timepoints after treatment. F-G: IL-6 cytokine secretion in T24 and J82 bladder cancer cells quantified by ELISA 24h post treatment with 50μM etoposide or 5mM hydroxyurea. H-J: HaCaT cells or A375 melanoma cells were transfected with 1μg/ml herring testing DNA, treated with 50μM etopside (Eto) or 5mM hydroxyurea (HU) for 6h, and the expression of IL-6 mRNA was quantified by qRT-PCR, normalised to GAPDH mRNA and expressed as fold change over respective mock treatments. K: HaCaT or A375 cells were exposed to ionizing radiation (IR, 10Gy), and IFN-β mRNA expression was quantified by qRT-PCR after recovery for 48h. Data from qRT-PCR experiments are shown as mean values of triplicate samples; error bars represent SD. Experiments were performed at least three times. Statistical significance was calculated using unpaired Student’s t-tests; * p<0.05, ** p<0.01, *** p<0.001.

**Supplementary Figure 2.**
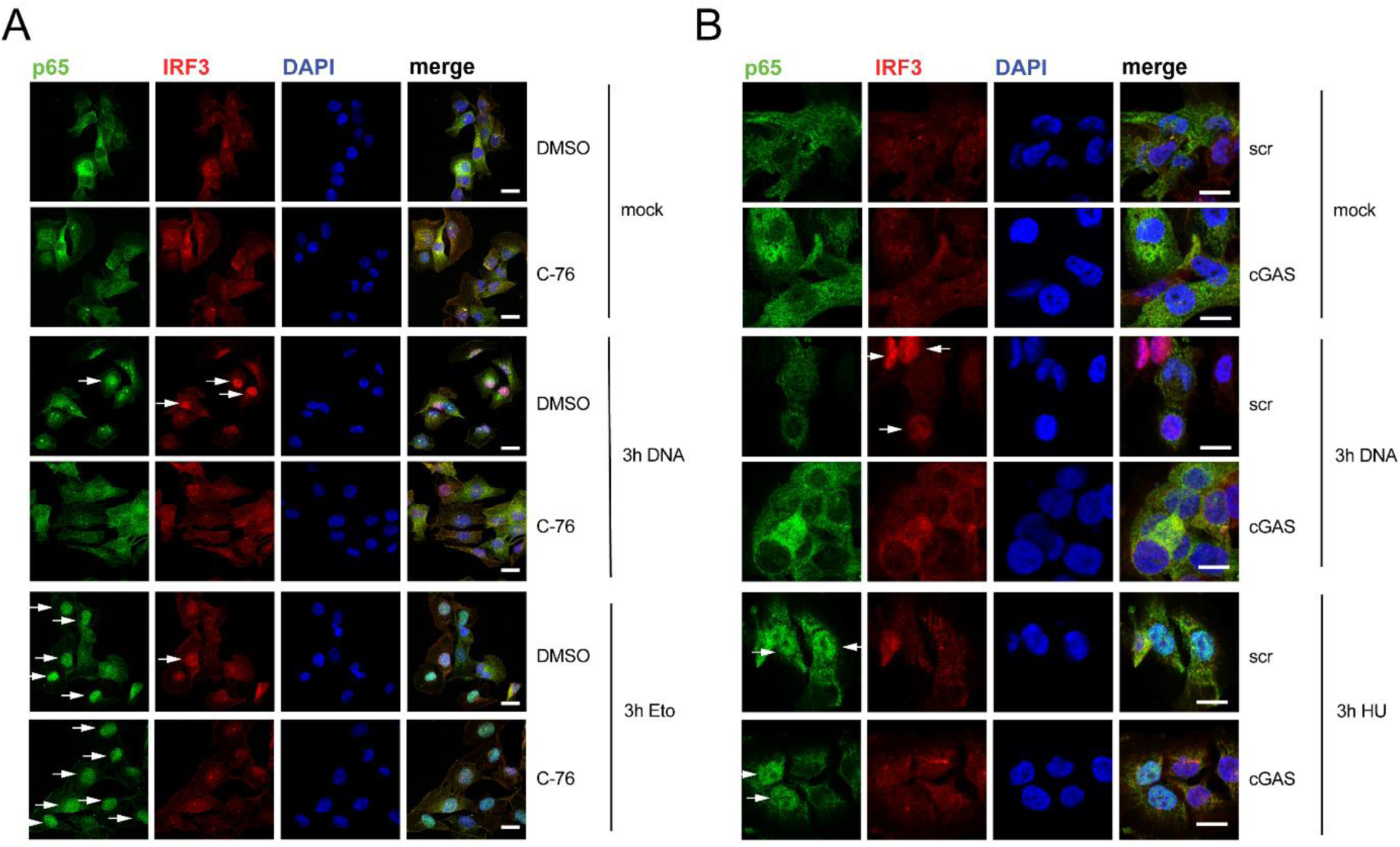
A: T24 bladder cancer cells were grown on coverslips and pre-treated with 50 μM cGAS inhibitor C-76 or DMSO control for 1h, prior to treatment with 50 μM etoposide (eto) or transfection with 5μg/ml herring testis DNA for 3h as indicated. Cells were fixed and stained for endogenous NF-κB p65 (green), IRF3 (red) and DNA (DAPI, blue). Scale bar: 20μm. B: T24 cells were grown on coverslips and transfected with a cGAS-targeting siRNA pool or scrambled siRNA (scr) for 48, and then treated with 5mM HU or transfected with 5μg/ml herring testis DNA for 3h. Cells were fixed and stained for endogenous NF-κB p65 (green), IRF3 (red) and DNA (DAPI, blue). Scale bar: 20μm.

**Supplementary Figure 3.**
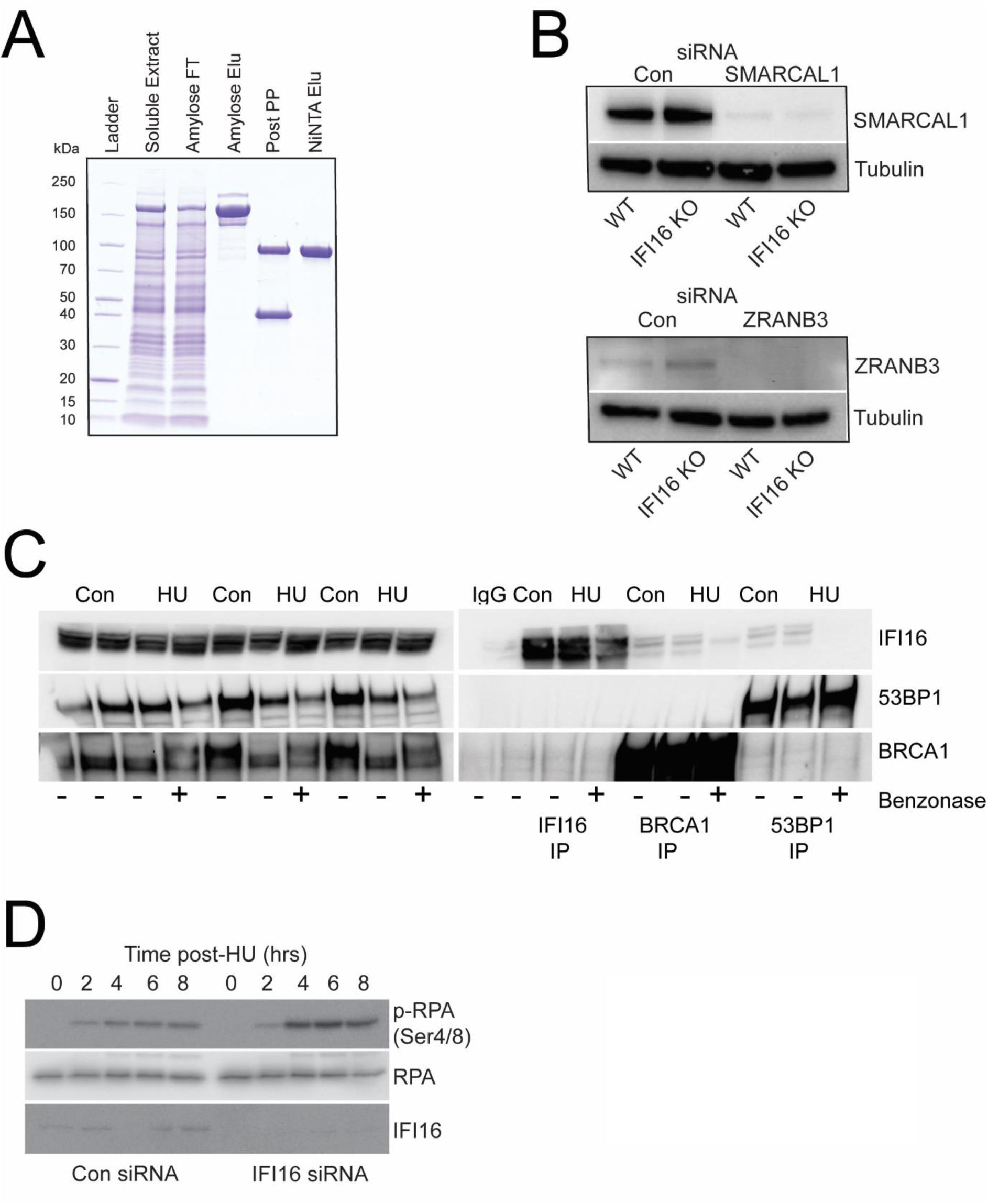
A: Purification of recombinant IFI16-His produced in *Sf9* cells. B: U2OS cells were transfected with the indicated siRNAs. After 48h cells were lysed and extracts resolved by western blotting following by probing with the indicated antibodies. C: HaCaT cells were treated with vehicle or 3 mM HU for 3 hr as indicated. Immunoprecipitation experiments were performed using Protein G beads and antibodies raised against BRCA1, 53BP1 or IFI16. IP eluates were resolved by western blotting and probing with the indicated antibodies. D: U2OS cells were transfected with a control siRNA or an siRNA targeting IFI16. After 48h cells were treated with 3 mM HU for the indicated times, then lysed and extracts resolved by western blotting followed by probing with the indicated antibodies.

